# Decoding Transcriptional Memory in Yeast Heat Shock and the Functional Implication of the RNA Binding Protein Mip6

**DOI:** 10.1101/2024.09.12.612644

**Authors:** Ana Tejada-Colón, Joan Serrano-Quílez, Carme Nuño-Cabanes, Susana Rodríguez-Navarro

**Affiliations:** Gene Expression and RNA Metabolism Laboratory, Instituto de Biomedicina de Valencia (CSIC). Jaume Roig, 11, 46010, Valencia, Spain

## Abstract

Cells not only adapt to environmental changes, but they also “remember” specific signals, allowing them to respond more rapidly to future stressors. This phenomenon, known as transcriptional memory, is orchestrated by a complex interplay of epigenetics, transcription regulators and RNA metabolism factors. Although transcriptional memory has been well-studied in various contexts, its role in the heat shock (HS) response of yeast remains largely unexplored. In our study, we delve into the dynamics of HS memory in wild-type yeast and a *mip6*Δ mutant, which lacks the RNA-binding protein Mip6. Notably, Mip6 has been shown to influence the expression of key stress-related genes and maintain low Msn2/4-dependent transcript levels under standard conditions. Our transcriptomic analysis offers novel insights into how yeast cells remember HS exposure. We uncover striking differences in gene expression patterns depending on whether genes are induced or repressed during HS memory. Furthermore, we find that an initial 15-minute heat shock triggers a response that becomes attenuated with just 5 additional minutes of stress. While the response kinetics between memory and non-memory conditions are similar, we report a subtle but important role for Mip6 in fine-tuning transcriptional memory and adaptation to heat stress. Biochemical and genetic evidence also suggests that Mip6 cooperates with alternative survival pathways independent of MSN2/4, and functionally interacts with the Rpd3 histone deacetylase complex, a key player in transcriptional memory and the HS response. These findings open up new avenues for understanding the molecular mechanisms behind heat stress adaptation in eukaryotes.

## Introduction

Cells have the ability to respond to environmental changes and adapt to adverse conditions such as high temperatures. Cells attempt to reduce and reverse proteotoxic damage and adapt to adverse environments through the activation of a series of mechanisms, including transcriptional changes, a global stop in translation and growth, and metabolic reprogramming. This results in the biosynthesis of protective molecules, such as molecular chaperones and trehalose, a disaccharide that serves a dual function by acting as a chemical chaperone and as an energy reserve that can be utilized during periods of cellular stress^1^. One of the most studied mechanisms to respond to high temperature is the Heat Shock Response (HSR), a specific transcriptional program that results in the overexpression of genes encoding stress-protective proteins, such as chaperones, and the silencing of genes related to cell growth^2–4^. The HSR is mainly regulated by the transcription factors Msn2/4 and Hsf1, which bind to regulatory elements of their target genes^5–8^.

The transcriptional reprogramming triggered by stress can be memorized, enabling cells to respond faster to future stimuli, a phenomenon known as “transcriptional memory”. This phenomenon can be observed for both stress-activated and stress-repressed genes and involves the activity of multiple players, including chromatin remodeling machinery, histone modifiers, transcription factors and even components of the mRNA degradation machinery^9–12^. Nuclear pores are also key elements in the establishment of transcriptional memory, serving as an anchoring structure for activators and repressors that function in transcriptional memory^13^. In addition, nuclear pores facilitate the formation of gene loops, a phenomenon associated with transcriptional memory that consists of the interaction between the promoter and the 3’ end region of the genes^14,15^.

In yeast, the study of transcriptional memory has focused on the response to two different stimuli: the lack of inositol in the medium, which provokes transcriptional activation of the gene *INO1*, and the shift of the carbon source in the medium from glucose to galactose, which results in the expression of the galactose metabolism genes^16–20^. In these studies, they report the role of the histone modifying complexes Rpd3L and Set1/COMPASS in the establishment of transcriptional repression memory (TREM), a process that regulates the rapid inactivation of silenced genes upon stress. At the beginning of transcription, the RNAPolII recruits the Set1/COMPASS complex, which methylates H3K4. Then, the Pho23 subunit of the Rpd3L complex binds H3K4met, triggering histone deacetylation by Rpd3L and the repression of the genes. In the absence of the Pho23 subunit, the cells do not present TREM, which suggest this mechanism is key for maintaining the transcriptional memory^16^.

We have recently demonstrated a functional role for the RNA-binding protein Mip6 in the yeast HSR. Under basal conditions, Mip6 localizes uniformly in the nucleus and cytoplasm, whereas upon heat shock it accumulates in stress granules and p-bodies, while changing the binding to specific mRNA targets^21^. Mip6 preferentially binds Msn2/4-dependent mRNAs under optimal conditions whereas, under HS, it binds preferentially to non-stress induced transcripts, such as genes encoding for ribosomal proteins^21^. Additionally, the absence of Mip6 results in the overexpression of many Msn2/4-dependent genes, including genes related with trehalose metabolism, but it provokes the underexpression of other genes, such as those encoding ribosomal proteins, suggesting a dual regulatory role for Mip6 in the expression of these genes^21,22^. Interestingly, we have also reported that Mip6 shows physical and genetic interactions with proteins that participate at different stages of mRNA synthesis and degradation processes, such as the RNAPolII subunit Rpb1, with proteins of the mRNA decay machinery Rrp6 and Xrn1, and the general mRNA export factor Mex67, all key players in the regulation of mRNA under stress and non-stress conditions^23–25^. However, the specific mechanism by which Mip6 can regulate specific targets depending on the environmental conditions needs to be further studied.

Using transcriptomic, genetic, and biochemical approaches, we investigate the changes that occur in yeast cells during the heat shock response (HSR) following either a single or two sequential heat shock (HS) events. Our study aims to determine the presence and significance of transcriptional memory in yeast cells in response to HS. Additionally, given Mip6’s known physical and functional interactions with the transcription machinery, we explore whether Mip6 plays a role in regulating transcriptional memory during heat shock.

## Results

### mRNA Abundance of Specific Genes Depends on Transcriptional Memory Upon Heat Shock Stress

The complete characterization of transcriptional memory induced by heat stress in yeast remains underexplored. To address this, we conducted a heat shock memory experiment using an overnight culture grown at 30 °C, which was divided into two flasks: one designated as “no memory”, refreshed with 30 °C medium, and the other as “memory” (Fig. 1). The “memory” flask was mixed with 51 °C-tempered medium to rapidly shift the temperature to 39 °C. This temperature was maintained for 20 min to induce the first heat shock (HS1), after which the flask was cooled to 30 °C for 60 min. Subsequently, the flask was subjected to a second heat shock at 39 °C for 20 min (HS2). The “no memory” flask underwent a single heat shock (39 °C for 20 min) after 80 min of post-refreshing. Samples for RT-qPCR analysis were collected at 0, 5, 10, 15, and 20 min during HS2 and HS’. This experiment was performed in triplicate with both WT and *mip6Δ* cells (Fig. 1).

**Fig. 1.**
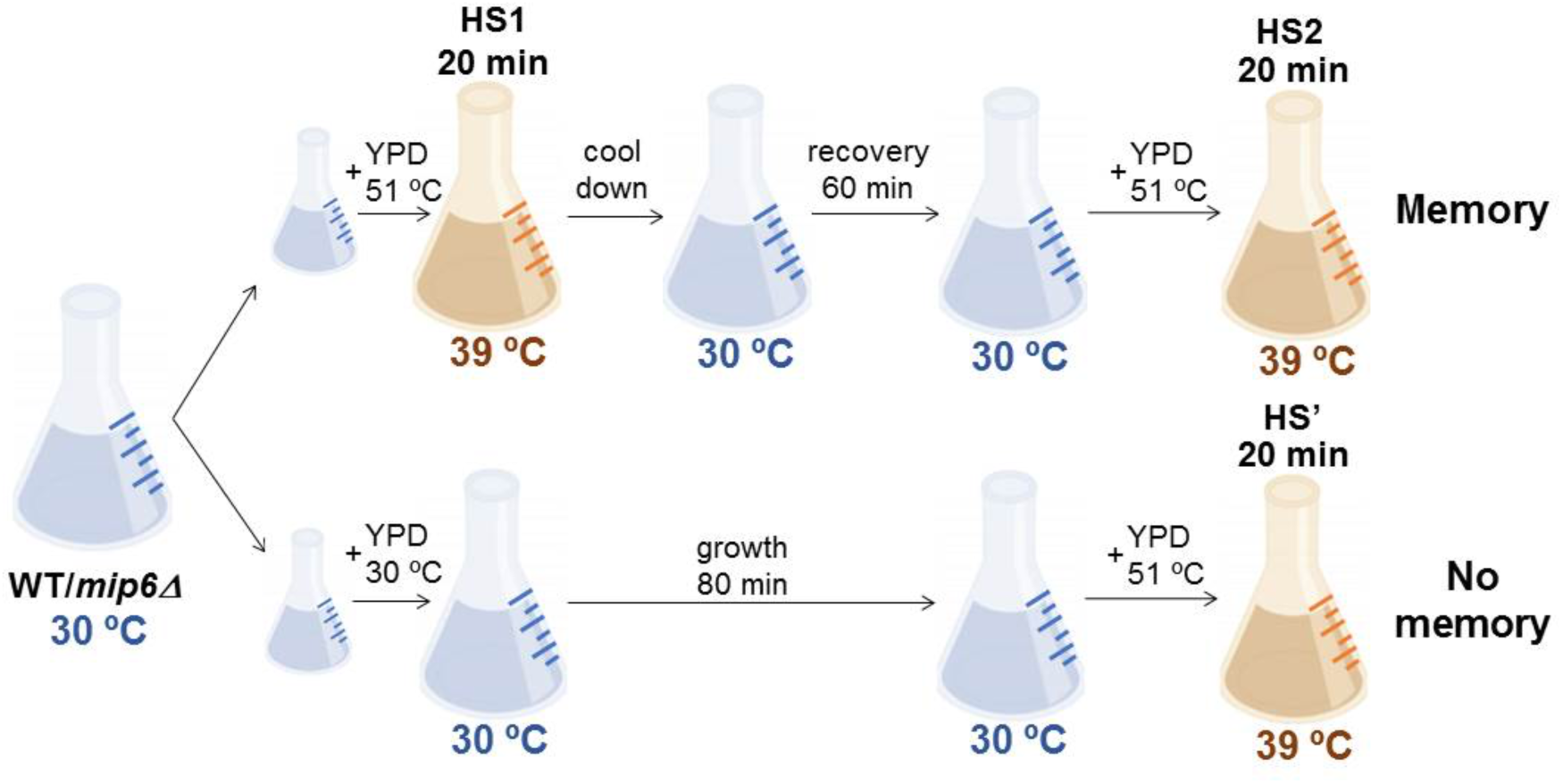
Experimental design for the study of the transcriptional memory associated with heat shock (HS). A cell culture was split into two flasks. One of them (memory) was exposed to a first 20 min heat shock and, after a 60 min recovery phase at optimal conditions, to a second 20 min heat shock. In the meantime, the other flask (no memory) was only exposed to a single 20 min heat shock. The heat shocks were performed by adding media at 51 °C.

We quantified specific heat shock response genes, using *SCR1* as a reference gene. In both WT and *mip6Δ* cells, the “no memory” condition resulted in a more robust activation of heat-responsive genes, including *HSP12*, *HSP78*, *HSP26*, *TPS1*, *TPS2*, and *FAA1*, as these transcripts reached higher levels more rapidly (Fig. 2a). The expression of *ACC1*, a heat shock-repressed gene, did not exhibit a differential pattern between memory and no memory conditions. When comparing WT and *mip6Δ* cells, no significant differences were detected. However, as expected, *mip6Δ* cells displayed higher transcript levels under heat shock, a phenotype observed under both memory (HS2) and no memory (HS’) conditions (Fig. 2a).

**Fig. 2.**
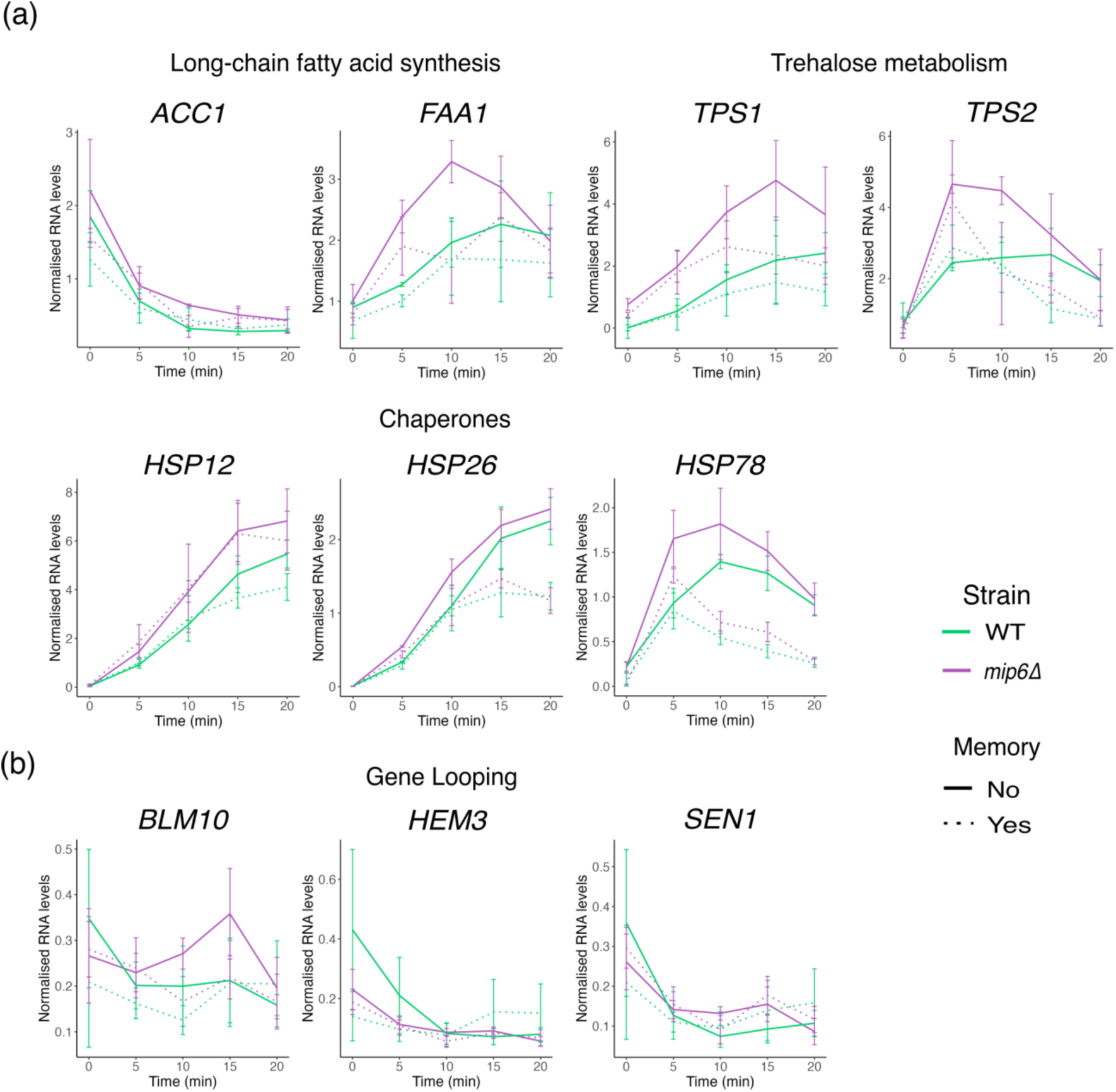
The “no memory” condition resulted in a higher activation of heat-responsive genes. (A) *ACC1, FAA1, TPS1*, *TPS2, HSP12*, *HSP78* and *HSP26* RNA levels measured by qRT-PCR experiments from WT and *mip6Δ* strains following the experiment previously described. Values were calculated by using ΔΔCt method. Data represent the mean and the standard error (SE) of three biological replicates. (B) *HEM3*, *SEN1*, and *BLM10* RNA levels measured by qRT-PCR experiments from WT and *mip6Δ* strains following the experiment previously described. Values were calculated by using ΔΔCt method. Data represent the mean and the standard error (SE) of three biological replicates.

To further explore the heat shock memory response, we measured the levels of three transcripts known to undergo gene looping, which is known to be regulated by transcriptional memory; *HEM3*, *SEN1*, and *BLM10*. Notably, in WT cells, *HEM3* and *SEN1* (both heat shock-repressed) showed a slight increase in the “no memory” condition compared to the “memory” condition, whereas *mip6Δ* lacks this phenotype (Fig. 2b). Additionally, *BLM10* expression was activated after 15 min of heat shock in *mip6Δ* cells, whereas WT cells maintained a flat or repressed expression. These findings suggest that Mip6 may play a role in gene looping, thereby influencing transcriptional memory.

We also investigated the expression of ribosomal protein (RP) genes, which are known to be downregulated under various stress conditions, including heat stress^26–28^. Notably, Mip6 has been reported to bind to RP gene transcripts during the heat shock response^21^. Our results showed that RP mRNA levels decreased similarly at 5 and 10 min post-heat shock (Fig. 3). However, in the “memory” samples, there was a tendency towards adaptation, with increased expression observed at the 20-minute time point, a response not seen in the “no memory” samples. Interestingly, in the *mip6Δ* mutant exhibited a different memory effect compared to the WT strain for several RP genes: *RPL6B*, *RPL23A*, *RPS10B*, *RPS17B*, and *RPS27A* (Fig. 3), suggesting a potential role for Mip6 in the regulation of transcriptional memory in these genes.

**Fig 3.**
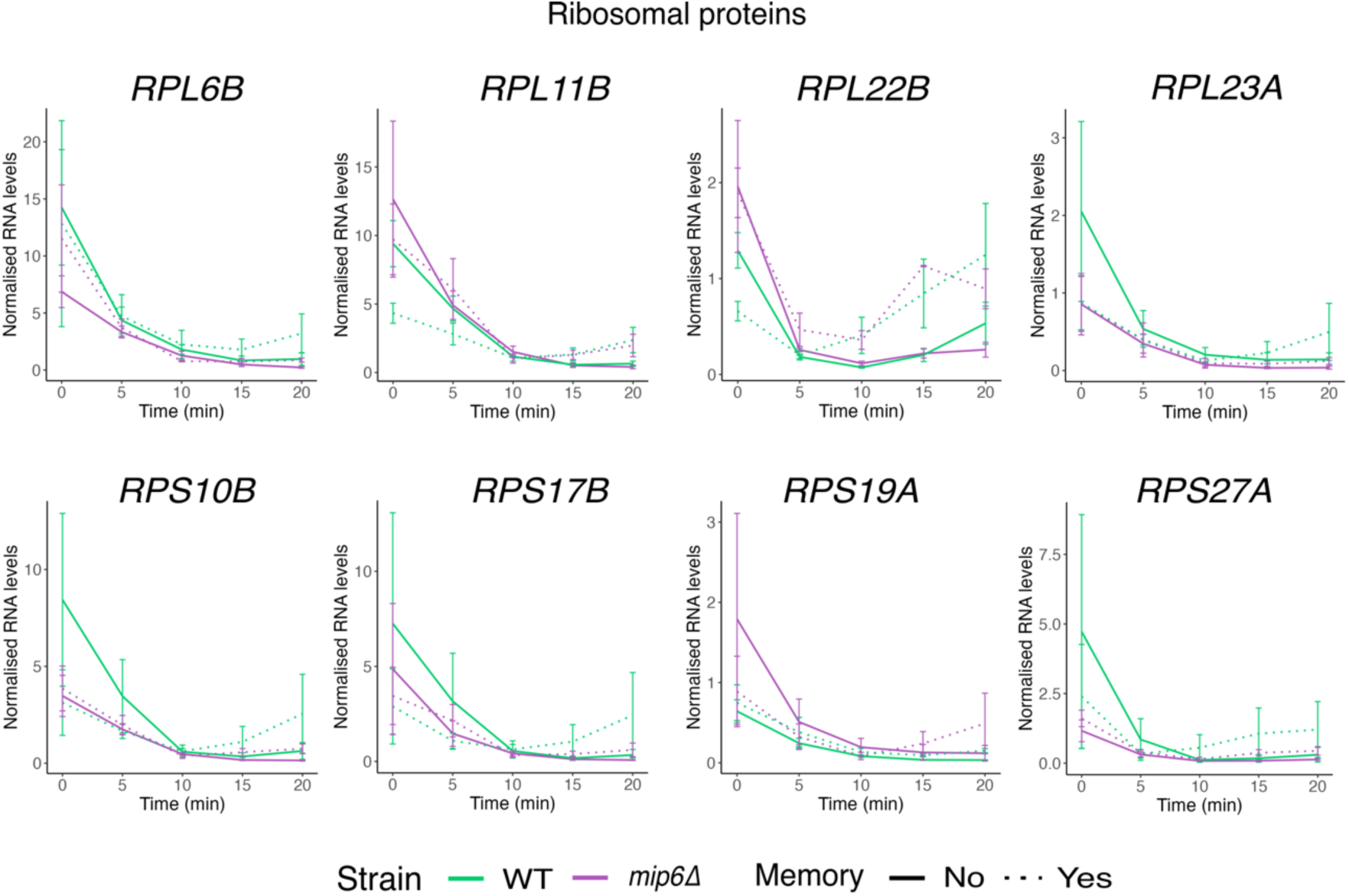
Ribosomal Protein (RP) genes are sensitive to transcriptional memory upon heat shock at 39 °C. *RPL6B, RPL11B, RPL22B RPL23A, RPS10B, RPS17B RPS19A* and *RPS27A* RNA levels measured by qRT-PCR experiments from WT and *mip6Δ* strains following the experiment previously described. Values were calculated by using ΔΔCt method and were normalized so that values of time 0 were equal to 1. Data represent the mean and the standard error (SE) of three biological replicates.

### Global Analysis of Transcriptional Memory Following Mild Heat Shock

To further investigate the changes in gene expression that occur when cells have been previously exposed to heat shock and to understand how the absence of Mip6 influences this mechanism, an RNA-Seq analysis was conducted, following the experimental design outlined above (Fig. 1). Samples were collected at 0, 15, and 20 min after the onset of heat stress.

As an initial overview of the RNA-Seq results, a principal component analysis (PCA) was performed to identify the primary variables accounting for the differences between the analyzed samples (Fig. 4a). The PCA provided a remarkably clear distinction between individual transcriptomes. Principal Component 1 (PC1), which is primarily dependent on temperature and time, accounted for 55.5 % of the variance, separating the samples based on growth conditions—optimal (0-min time point) versus stress (15-min and 20-min time points). Thus, more than half of the observed differences were attributable to the environmental conditions in which the cells were grown. Principal Component 2 (PC2), representing 16.3 % of the variance, clearly distinguished the samples based on whether the cells had been previously exposed to heat stress (“memory”) or not (“no memory”). This suggests that more than one-fifth of the differences were due to transcriptional memory. Consequently, other potential variables, such as strain differences or biological replicates, played a less significant role in explaining the observed variance, as nearly 75 % of the differences were attributed to stress and memory effects.

**Fig. 4.**
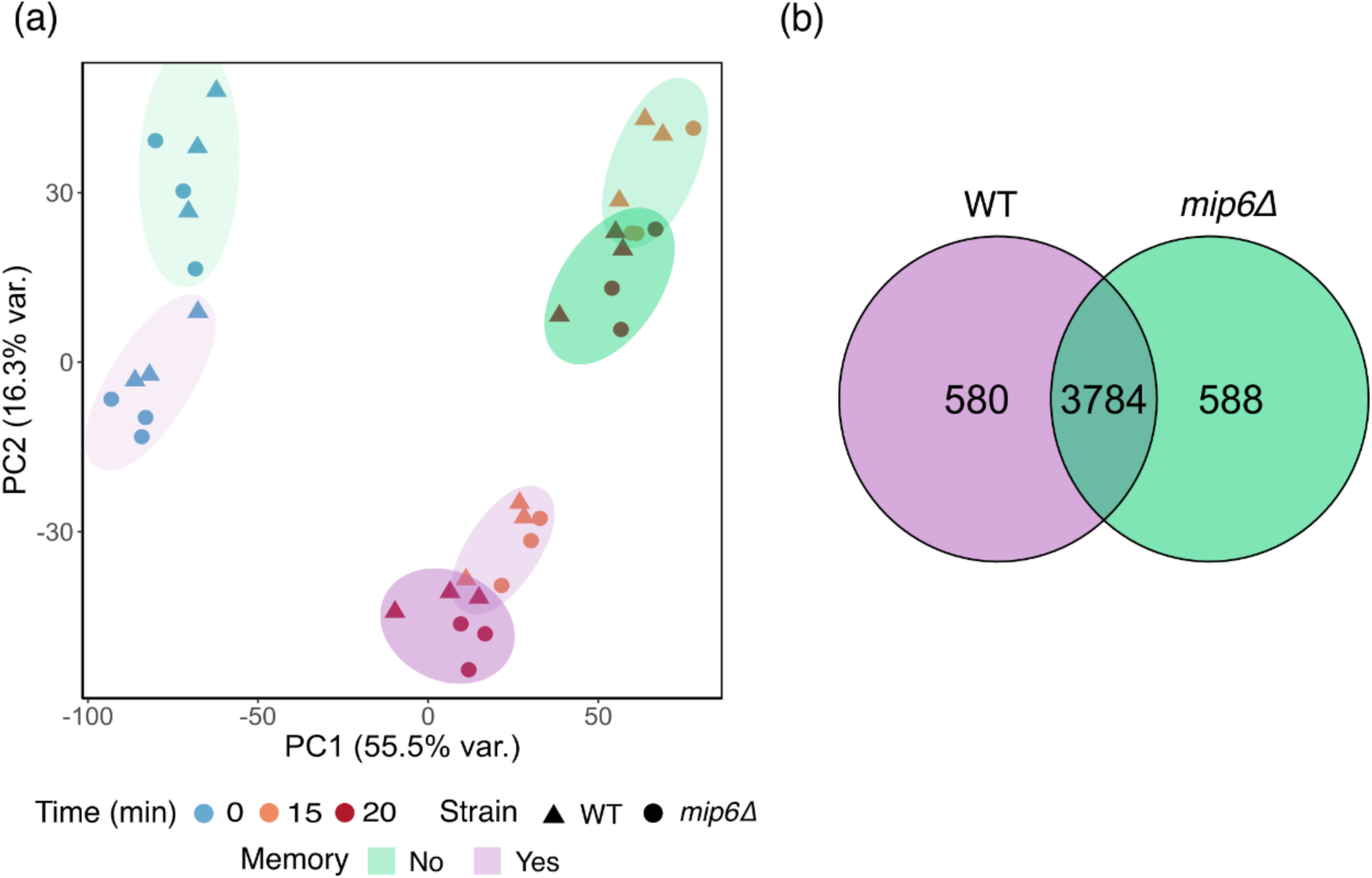
Heat shock stress response is different when yeast cells have already suffered a previous heat exposition. (A) PCA score plot from the RNA-Seq analysis of the WT and *mip6Δ* collected at different times during a heat shock treatment (0, 15 or 20 min at 39 °C). The samples were previously exposed to a 20 min heat shock at 39 °C with a 60 min recuperation period at 30 °C (memory) or were previously grow only under optimal conditions (no memory). Three biological replicates were carried out. (B) Venn diagram of genes significantly affected by transcriptional memory to heat shock in WT and *mip6Δ* strains, as obtained by maSigPro.

In both WT and *mip6Δ* strains, more than two-thirds of the yeast genes exhibited differential expression during the HSR depending on prior exposure to heat stress: 4364 genes in the WT strain and 4372 in the *mip6Δ* mutant out of a total of 6172 (Fig. 4b). Although most genes that exhibit transcriptional memory in the mutant also display this memory in the WT strain, there are 580 genes with a unique differential response in WT and 588 genes in the mutant (Fig. 4b). These differences suggest a potential role for Mip6 in regulating transcriptional memory, as the absence of this protein causes some genes to lose their memory-associated expression patterns.

To identify differences between WT and *mip6Δ*, we further analyzed the set of genes exhibiting unique behavior in the mutant. Out of the 588 genes, significantly overrepresented Gene Ontology (GO) terms related to biological processes included sucrose catabolic process (GO:0005987) and oligosaccharide metabolic process (GO:0009311), maltose metabolic process (GO:0000023), and disaccharide metabolic process (GO:0005984). This is interesting, since MSN2/4 are key regulators of the metabolic branch of the HSR ^28^ and Mip6 interacts with them^21,22^.

### Mip6 Cooperates with Different Survival and Adaptation Mechanisms Independent of MSN2/4

Recent studies have revealed a combination of survival strategies that collectively protect essential proteins in yeast^28^. These strategies rely on *MSN2/4* and/or *HSF1*. Msn2/4 broadly reprograms transcription, triggering responses to oxidative stress, and promotes the biosynthesis of the protective sugar trehalose and glycolytic enzymes. In contrast, Hsf1 primarily induces the synthesis of molecular chaperones and modulates the transcriptional response during prolonged mild heat stress (adaptation). The proposed model suggests that the deletion of Msn2/4 negatively impacts the metabolic branch of the HSR, while Hsf1 is crucial for the efficient upregulation of the chaperone branch. Additionally, the model highlights the involvement of other factors in regulating survival and adaptation. Notably, in the absence of Hsf1 and Msn2/4, several genes involved in sporulation were still upregulated under stress, indicating the existence of an emergency escape program independent of Hsf1 and Msn2/4^28^. Mip6 is functionally linked to sporulation^29^ and current research in our lab suggest its participation in early meiotic gene (EGM) regulation (not shown). We closely examined our differentially expressed genes (DEGs) list and filtered it for terms related to sporulation, meiosis, and mating. We identified at least 27 genes associated with these processes that were affected specifically by the absence of Mip6 and 31 affected in the WT (Table S1). Interestingly, some of these genes show a pattern that fits with a role for Mip6 as a repressor or are specific for meiosis recombination (*GIP1, HOP2, MEI5, SAE3, REC107, RIM4*). To functionally assess this non-redundant role between Msn2/4 and an alternative factor that might contribute to the alternative scape program under HS, we constructed double and triple mutants to compare the growth of WT, *mip6Δ, msn2Δ, msn2Δmip6Δ, msn4Δ, msn4Δmip6Δ, msn2Δmsn4Δ*, and *msn2Δmsn4Δmip6Δ* strains. Both the *MSN2* and *MSN4* single deletions led to growth defects under heat stress, underscoring the importance of these factors in heat stress survival (Fig. 5a). Unexpectedly, when both deletions were combined, the cells grew at the same rate as the WT strain in our experiments (Fig. 5a). In contrast, no significant genetic interaction with *MIP6* was observed under heat stress conditions. However, we observed a non-statistically significant trend in the *msn2Δmip6Δ* strain, which showed a slight increase in growth rate compared to the *msn2Δ* single mutant, a trend also observed in the triple mutant (Fig. 5a).

**Fig. 5.**
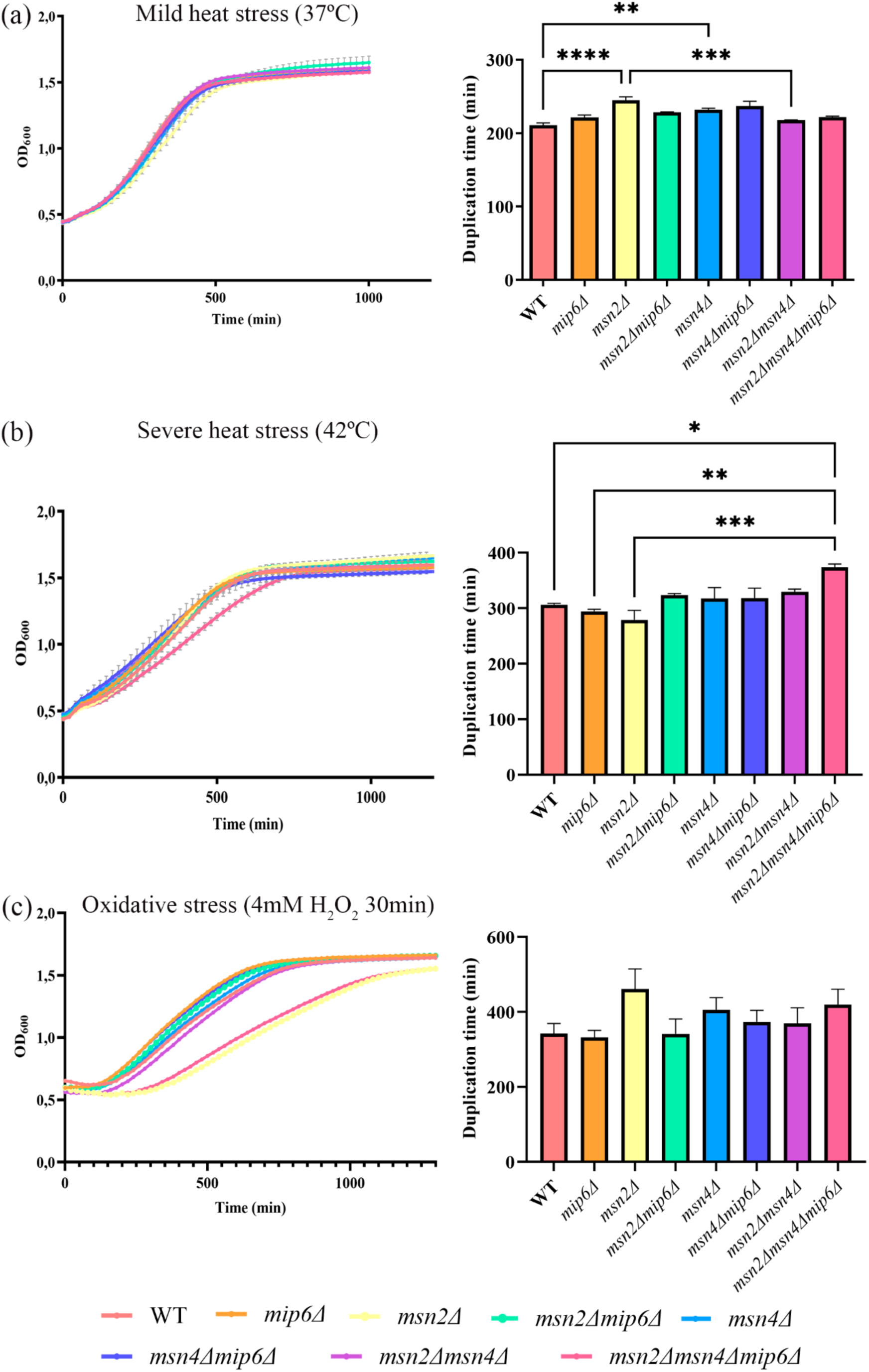
*MIP6, MSN2 and MSN4* genetically interact upon severe heat and oxidative stress. (A) Growth curve of wild-type (WT), *mip6Δ, msn2Δ, msn2Δmip6Δ, msn4Δ, msn4Δmip6Δ, msn2Δmsn4Δ* and *msn2Δmsn4Δmip6Δ* strains upon mild heat shock (37 °C). Data represent the mean and the standard error (SE) of three biological replicates (left). Duplication time of wild-type (WT), *mip6Δ, msn2Δ, msn2Δmip6Δ, msn4Δ, msn4Δmip6Δ, msn2Δmsn4Δ* and *msn2Δmsn4Δmip6Δ* strains upon mild heat shock (37 °C). Data represent the mean and the standard error (SE) of three biological replicates (right). (B) Growth curve of wild-type (WT), *mip6Δ, msn2Δ, msn2Δmip6Δ, msn4Δ, msn4Δmip6Δ, msn2Δmsn4Δ* and *msn2Δmsn4Δmip6Δ* strains upon severe heat shock (42 °C). One representative replicate is shown (left). Duplication time of wild-type (WT), *mip6Δ, msn2Δ, msn2Δmip6Δ, msn4Δ, msn4Δmip6Δ, msn2Δmsn4Δ* and *msn2Δmsn4Δmip6Δ* strains upon mild heat shock (37 °C). Data represent the mean and the standard error (SE) of three biological replicates (right). (C) Growth curve of wild-type (WT), *mip6Δ, msn2Δ, msn2Δmip6Δ, msn4Δ, msn4Δmip6Δ, msn2Δmsn4Δ* and *msn2Δmsn4Δmip6Δ* strains after an oxidative shock (4 mM H2O2 30 min). One representative replicate is shown (left). Duplication time of wild-type (WT), *mip6Δ, msn2Δ, msn2Δmip6Δ, msn4Δ, msn4Δmip6Δ, msn2Δmsn4Δ* and *msn2Δmsn4Δmip6Δ* strains after an oxidative shock (4 mM H2O2 30 min). Data represent the mean and the standard error (SE) of six biological replicates (right).

The trend observed in the triple mutant prompted us to investigate the genetic interaction between *MIP6, MSN2*, and *MSN4* under more severe stress conditions, such as growth at 42°C. The *msn2Δmsn4Δmip6Δ* mutant exhibited a growth defect compared not only to the WT strain but also to the corresponding single mutants (Fig. 5b), indicating a genetic interaction between *MSN2, MSN4* and *MIP6* under severe heat shock. Moreover, in this context, the absence of only one transcription factor did not significantly impact cell survival, suggesting that the presence of either *MSN2* or *MSN4* is sufficient to trigger an effective HSR.

Msn2/4 also regulate other types of stress responses, such as oxidative stress^30,31^. Additionally, Mip6 appears to play a role in fatty acid oxidation in peroxisomes and in maintaining redox balance^32^. Considering all these findings, Mip6 might also participate in the oxidative stress response (OxSR), which is closely linked to the HSR^3^. To explore this possibility, we examined the genetic interaction between *MIP6, MSN2* and *MSN4* under oxidative stress by performing growth assays on WT, *mip6Δ, msn2Δ, msn2Δmip6Δ, msn4Δ, msn4Δmip6Δ, msn2Δmsn4Δ,* and *msn2Δmsn4Δmip6Δ* strains following oxidative shock with 4 mM H2O2 for 30 min. Although not statistically significant, differences in the duplication time of the strains were observed following oxidative shock (Fig. 5c). The *msn2Δ* mutant displayed a growth defect that was mitigated by the deletion of either *MIP6* or *MSN4*. However, the triple mutant exhibited the poorest growth among all combinations (Fig. 5c). These results indicate a genetic interaction between *MIP6, MSN2*, and *MSN4* under oxidative stress.

In summary, the *msn2Δmsn4Δmip6Δ* triple mutant is the only strain to exhibit a pronounced growth defect under severe heat shock (Fig. 5b). This finding not only demonstrates a genetic interaction between *MSN2, MSN4*, and *MIP6* under severe heat shock but also suggests that these proteins participate in distinct cell survival mechanisms.

### Mip6 Functionally Interacts with Histone Deacetylase Rpd3

The analysis of overrepresented GO terms among genes affected by memory in *mip6Δ* but not in the WT strain included categories related to sugar metabolism. Furthermore, we conducted a comprehensive analysis of the genes significantly affected by memory. The analysis revealed that the differential response to memory in both strains encompasses genes encoding proteins involved in regulating gene expression through histone modification, such as various subunits belonging to the SAGA and PAF1 complexes. However, notably, only in the *mip61′* mutant did we identify three genes belonging to the Rpd3L deacetylase complex (*ASH1*, *DOT1*, *HOS2*), and a gene from the Rpd3 small (Rpd3S) complex (*EAF3*) that were affected by memory. Significantly, among the factors that co-purified with Mip6-TAP in our tandem affinity purification (TAP) experiments, we identified Rpd3 and Gcn5 (unpublished data).

Rpd3 is behind transcriptional memory mechanism in yeast^16^. Thus, we find very interesting the possible functional connection between Mip6 and Rpd3. To confirm the physical interaction between Mip6 and Rpd3, we performed a co-immunoprecipitation assay by expressing Mip6-GFP from a plasmid in an Rpd3-TAP *mip6Δ* strain. The experiment was conducted under optimal conditions, following heat shock (39°C for 20 min), and with impaired Mip6-Mex67 interaction using the Mip6W442A mutant. As shown in Fig. 6a, Mip6 physically co-purifies with the histone deacetylase Rpd3. This interaction is maintained under both optimal and heat stress conditions and is independent of the Mip6-Mex67 interaction, as the Mip6W442A mutation does not disrupt the co-purification (Fig. 6a). Thus, Mip6 interacts with Rpd3 in a Mex67-independent manner under both optimal and heat stress conditions.

**Fig. 6.**
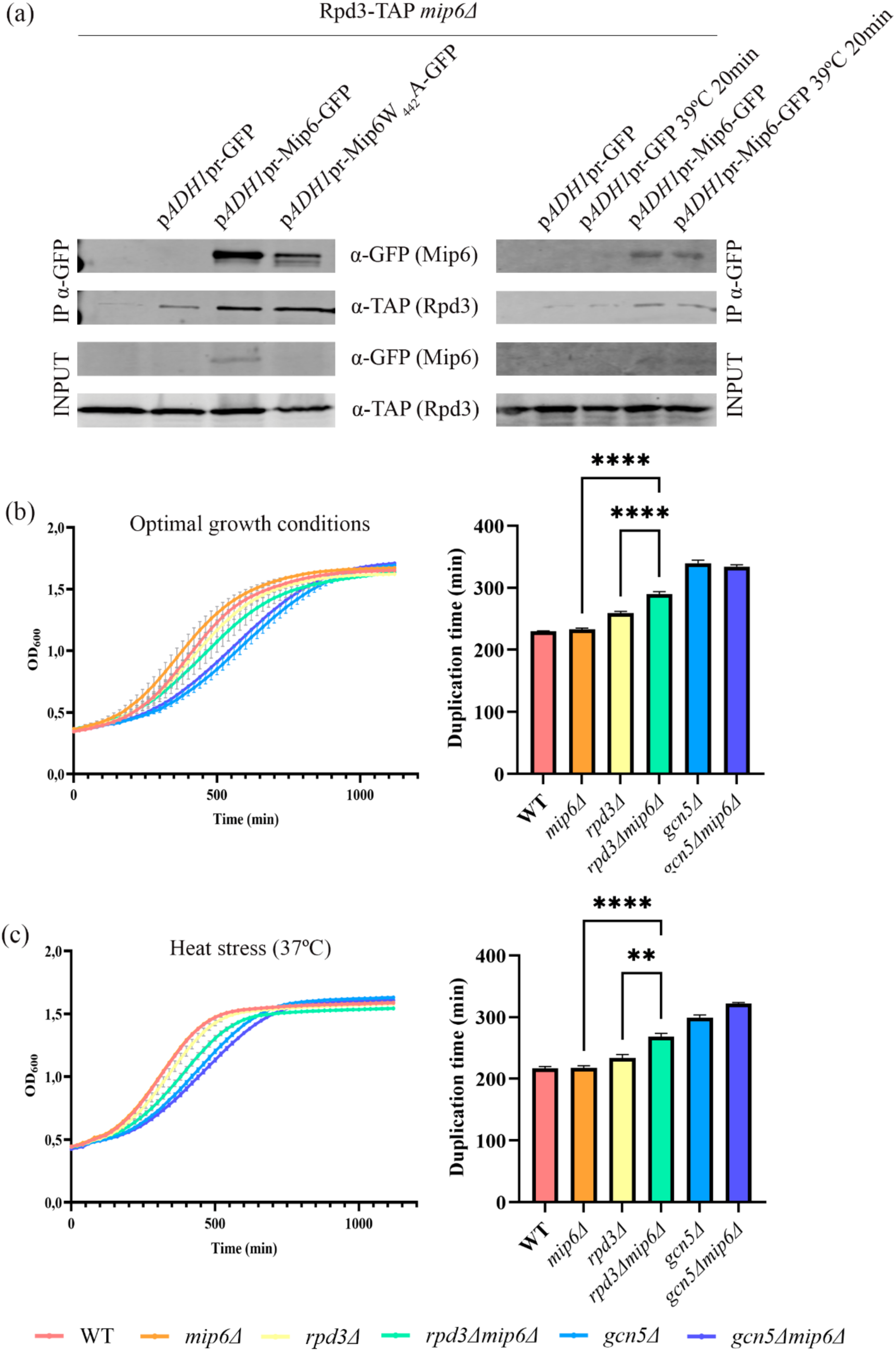
Mip6 physically, genetically and functionally interacts with HDAC Rpd3. (A) Immunoprecipitation of Mip6-GFP or Mip6W442A-GFP expressed from plasmid in Rpd3-TAP *mip6Δ* strain under optimal conditions or following a heat shock at 39 °C for 20 min. An empty plasmid and a strain that expresses no plasmid were used as negative controls. Mip6-GFP and Rpd3-TAP were detected by Western blotting using the indicated antibodies. (B) Growth curve of WT, *mip6Δ, rpd3Δ, rpd3Δmip6Δ, gcn5Δ* and *gcn5Δmip6Δ* strains under optimal conditions. Data represent the mean and the standard error (SE) of three biological replicates (left). Duplication time of WT, *mip6Δ, rpd3Δ, rpd3Δmip6Δ, gcn5Δ,* and *gcn5Δmip6Δ* strains under optimal conditions. Data represent the mean and the standard error (SE) of three biological replicates. **** p value < 0.001 (right). (C) Growth curve of WT, *mip6Δ, rpd3Δ, rpd3Δmip6Δ, gcn5Δ* and *gcn5Δmip6Δ* strains upon heat stress (37 °C). Data represent the mean and the standard error (SE) of three biological replicates (left). Duplication time of WT, *mip6Δ, rpd3Δ, rpd3Δmip6Δ, gcn5Δ,* and *gcn5Δmip6Δ* strains upon heat stress (37 °C). Data represent the mean and the standard error (SE) of three biological replicates. **** p value < 0.001; ** p value < 0.01 (right).

We next examined the growth of WT, *mip6Δ, rpd3Δ, rpd3Δmip6Δ, gcn5Δ*, and *gcn5Δmip6Δ* strains. The growth assay was conducted under both optimal conditions and heat stress (37 °C) to compare growth curves and duplication times. As illustrated in Fig. 6b-c, the *rpd3Δmip6Δ* double mutant exhibited poorer growth compared to the corresponding single mutants, confirming a negative genetic interaction between *MIP6* and *RPD3*. In contrast, no significant interaction was observed for *GCN5* when deleted in the absence of *MIP6* (Fig. 6b-c).

The genetic and physical interaction between Mip6 and Rpd3 may also impact the expression levels of Msn2/4-regulated transcripts, as the absence of either protein individually affects their expression^21,33^. To explore this, we analyzed the mRNA levels of *HSP12* and *CTT1*, two Msn2/4-dependent transcripts, in WT, *mip6Δ, rpd3Δ*, and *rpd3Δmip6Δ* strains by qPCR. In contrast to the increase observed with *MIP6* deletion (Fig. 7a), both *HSP12* and *CTT1* mRNA levels decreased in the absence of Rpd3 (Fig. 7b). Furthermore, the double mutant *rpd3Δmip6Δ* displayed mRNA levels similar to the *rpd3Δ* single mutant (Fig. 7b). This suggests that both proteins may function in the same pathway, with Rpd3 being essential for Mip6’s role in this process.

**Fig. 7.**
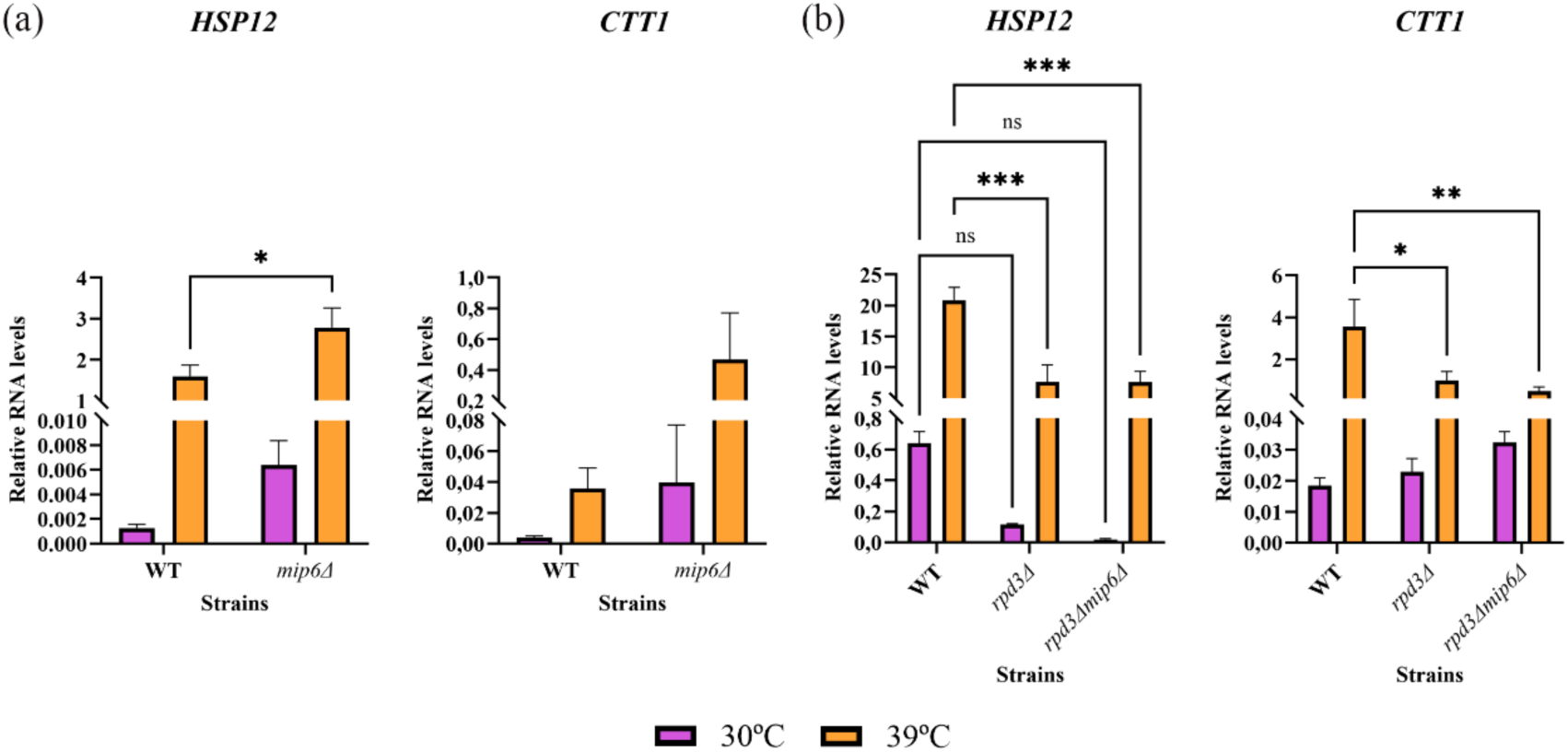
Analysis of the synergistic effect of HDAC deletion with *mip6Δ* in the expression levels of *HSP12* and *CTT1*. (A) *HSP12* and *CTT1* RNA levels measured by qRT-PCR experiments from WT and *mip6Δ* strains. Total RNA was obtained from yeast cultures under optimal conditions and after a heat shock of 20 min at 39 °C. Values were calculated by using ΔΔCt method. Data represent the mean and the standard error (SE) of three biological replicates. * p value < 0.05. (B) *HSP12* and *CTT1* RNA levels measured by qRT-PCR experiments from WT, *rpd3Δ* and *rpd3Δmip6Δ* strains. Total RNA was obtained from yeast cultures under optimal conditions and after a heat shock of 20 min at 39 °C. Values were calculated by using ΔΔCt method. Data represent the mean and the standard error (SE) of three biological replicates. *** p value < 0.005; ** p value < 0.01; * p value < 0.05.

### Memory amplifies the expression of specific genes while attenuating the response of others

To globally analyze how memory affects gene response, we grouped all significant genes impacted by memory into four different clusters, based on the differential expression analysis from MaSigPro and the expression pattern of each gene (Fig. 8). In genes with an increased expression during HS, we observed that memory has a wider impact in negatively affecting the response of 1738 genes (cluster 1) while the positive influence in the expression is limited to 497 genes (cluster 3). The mutant exhibits a behavior similar to that of the WT, although differences are shown by heatmaps of the clusters on specific gene-sets (Fig. S1). Thus, in terms of number of genes, expression of most yeast genes is negatively affected by memory. Genes whose expression is repressed during the HSR are shown in clusters 2 and 4 (Fig. 8 and Fig. S1). In contrast to induced genes, repression is always higher in the “no memory” samples than in the “memory” ones, either because they already present lower expression levels under optimal conditions (cluster 2) or because, even if their optimal mRNA levels are similar in both samples, their reduction upon heat shock softens with memory (cluster 4). With these clusters, we performed a gene overrepresentation test for biological processes (Table 1).

**Fig. 8.**
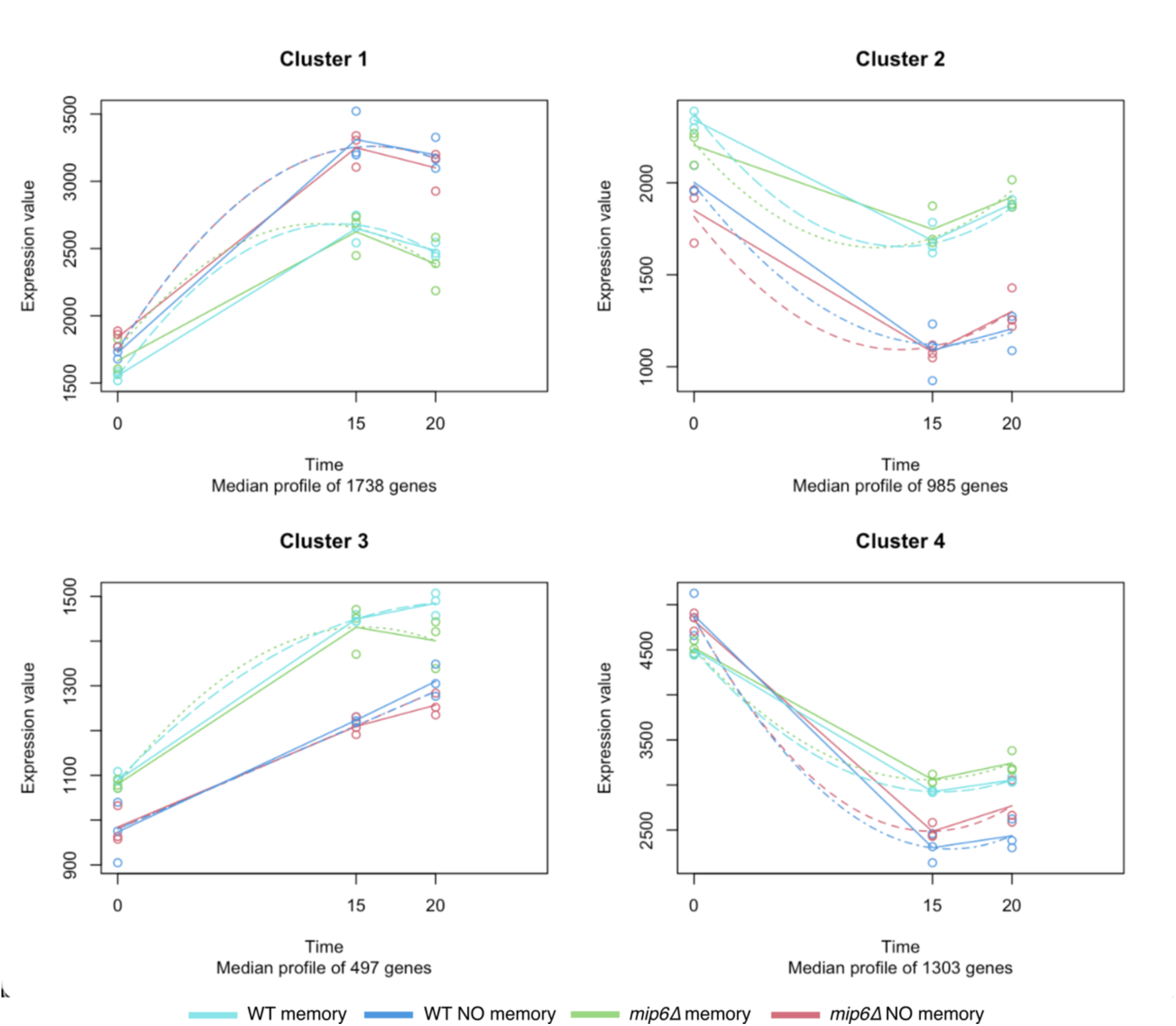
Clustering of the genes that present differential expression between memory and no memory in WT and *mip6Δ* strains. Genes with differential expression between “memory” (light blue) and “no memory” (dark blue) in WT and between “memory” (green) and “no memory” (red) in *mip6Δ* were plotted in four different clusters according to their expression profile along the heat shock response. Data represent the mean value of three biological replicates.

**Table 1.** Gene ontology analysis of DE genes. Table containing the overrepresented GOs in each of the maSigPro clusters.

|  |  |
| --- | --- |
| **Cluster 1** | monosaccharide catabolic process (GO:0046365) |
|  | aerobic electron transport chain (GO:0019646) |
|  | trehalose metabolic process (GO:0005991) |
|  | ubiquitin-dependent protein catabolic process (GO:0006511) |
|  | negative regulation of gluconeogenesis (GO:0045721) |
|  | ketone catabolic process (GO:0042182) |
|  | aldehyde catabolic process (GO:0046185) |
|  | fatty acid catabolic process (GO:0009062) |
| **Cluster 2** | tRNA metabolic process |
|  | poly(A)-dependent snoRNA 3'-end processing (GO:0071051) |
|  | ribosomal large subunit assembly (GO:0000027) |
|  | tRNA transcription by RNA polymerase III (GO:0042797) |
|  | ribosomal small subunit biogenesis (GO:0042274) |
|  | mRNA catabolic process (GO:0006402) |
|  | nucleolar large rRNA transcription by RNA polymerase I (GO:0042790) |
| **Cluster 3** | mitochondrial translation (GO:0032543) |
| **Cluster 4** | biogenic amine biosynthetic process (GO:0042401) |
|  | amino acid metabolic process (GO:0006520) |
|  | protein N-linked glycosylation (GO:0006487) |
|  | ergosterol biosynthetic process (GO:0006696) |
|  | ribonucleoside monophosphate metabolic process (GO:0009161) |

Among genes that are induced during HS (clusters 1 and 3), cluster 1 is more functionally heterogeneous, being most of the overrepresented in terms related to the obtention of energy, either by biomolecule catabolism or by respiration. It is worth noting that this cluster, which is less induced when memory is applied, contains the majority of genes related to trehalose metabolism. This might indicate that the previous exposure to a HS has already provoked metabolic changes that are necessary for a correct response, so no further expression of these genes is necessary. On the other hand, in cluster 3, representing those that have a greater response with memory, the only overrepresented term relates to genes that are found in mitochondria.

Regarding HS-repressed genes, most GOs that appear in both clusters 2 and 4 are related to the reduced protein synthesis that occurs under these conditions. Whereas cluster 2 contains most genes related to the process of translation itself, cluster 4 encompasses genes involved in amino acid biosynthesis. The fact all these genes are less repressed when cells have been previously subjected to HS, might indicate that cells have already adapted these machineries to a lower activity level.

To compare the proportion of genes that change during the HS in memory and no memory, we represented these responses in Sankey plots (Fig. 9). From the total of 6172 analyzed genes, 1706 in the WT and 1615 in the mutant showed no induction or repression after 15 min of stress in the no memory samples, whereas in memory samples, 2119 genes in the WT and 1886 genes in the mutant showed this flat behavior. Interestingly, this soothing effect of memory in gene expression was observed through the strong activated and repressed genes (in dark green and red, respectively), which are reduced from 772 to 637 genes in the WT and from 876 to 797 genes in the *mip61′* mutant. A similar effect was also observed when comparing the time 0 with time 20 min.

**Fig 9.**
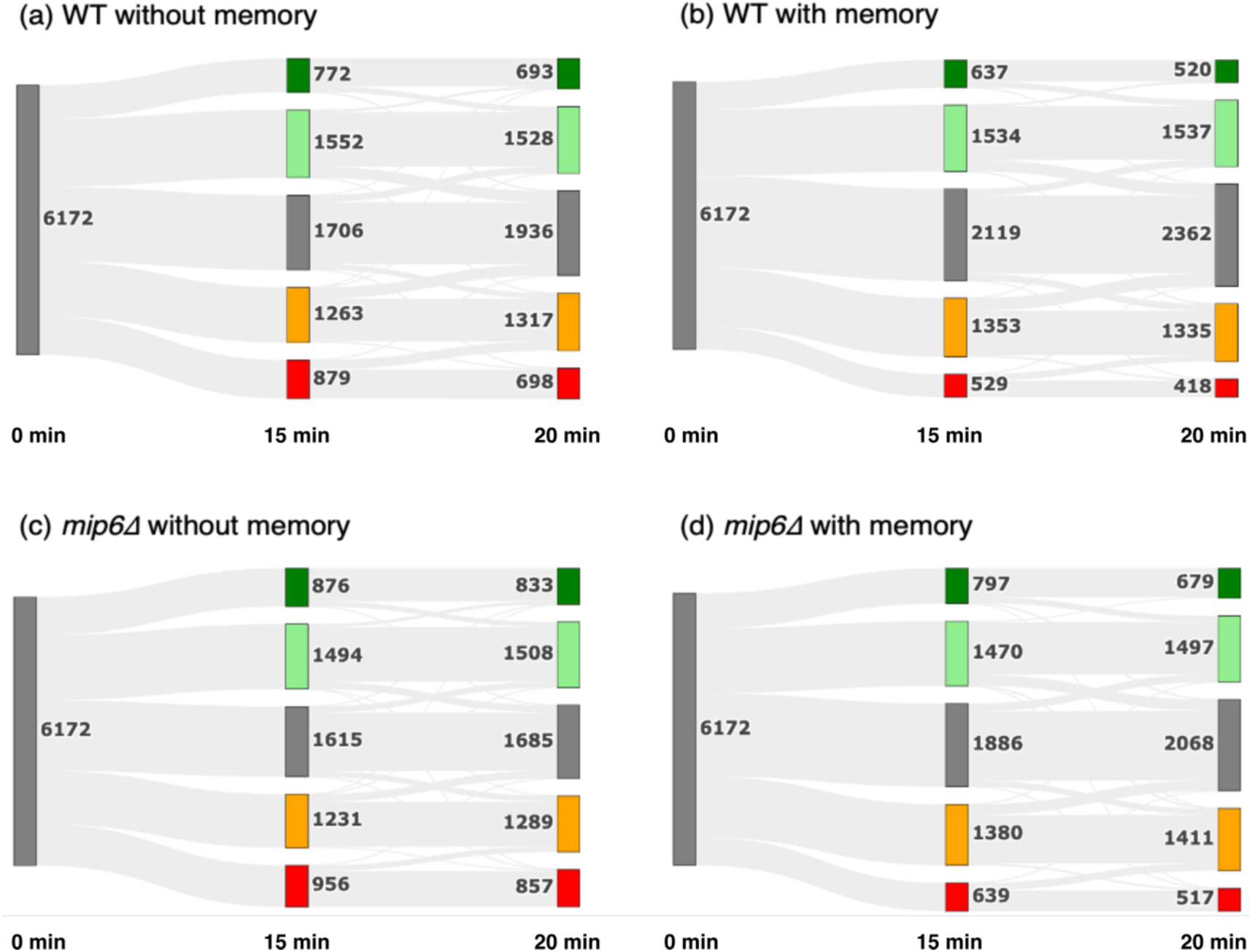
Sankey plots representing changes in gene expression along time under heat-stress. Genes are classified according to their expression level when compared to the initial time in the same strain and condition. In dark green, genes with a foldchange higher than 2. In light green, genes displaying a foldchange between 1.2 and 2. In gray, those genes with a foldchange lower than 1.2 and higher than −1.2 or not significantly affected. In orange, genes with a fold change between −1.2 and −2. In red, those with a foldchange lower than −2.

Then we analyzed the behavior of *mip6Δ* (Fig. 9). Absence of Mip6 changes the magnitude of the transcriptional adjustment increasing these DEGs at 15 min in 91 genes. Interestingly, the mutant showed a higher number of genes that are strongly activated and repressed (dark green and red, respectively) at both time points and conditions (memory and no memory), which reinforces Mip6 participation in fine-tuning their mRNA levels.

## Discussion

### Decoding transcriptional memory upon HS

Yeast cells exhibit a transcriptional memory mechanism, allowing them to respond more quickly to heat stress if they have been previously exposed to it. This memory can persist for up to 14 generations^12,18,19^ and has been documented in other organisms like Arabidopsis^34,35^. In yeast, most studies have focused on specific genes like galactose metabolism and *INO1* induction^18–20^. Exposure to heat stress in *S. cerevisiae* alters the expression of a significant portion of the genome, with more than two-thirds of genes showing modified expression upon a second exposure. Three distinct gene expression patterns were identified in our study:

1. Weakened Repression: About 2288 genes that are typically repressed during heat stress (clusters 2 and 4), show a reduced level of repression during a second exposure. This might be due to a faster adaptation to heat stress or the continued sequestration of these transcripts in stress granules and processing bodies. Interestingly, memory does not enhance repression in a significant number of genes
2. Reduced Activation: A subset of 1738 genes shows lower activation levels during the second heat exposure compared to the first. This may be due to incomplete recovery from the initial stress or a form of transcriptional memory, where proteins like Hsp12 are already upregulated, allowing a quicker response without needing to increase mRNA levels as much.
3. Enhanced Activation: Another set of 497 genes shows faster and stronger activation upon repeated heat exposure, reflecting the classical model of transcriptional memory. These genes include those involved in mitochondrial translation.

Mip6 is a RBP that regulates gene expression of certain stress-response genes; although other kind of target genes were also described, such as trehalose biosynthesis genes or ribosomal protein genes. Nevertheless, the exact mechanism by which this protein is able to regulate mRNA levels of these genes is still unknown, even if it has been related with different steps of the gene expression process. Thus, in this work, we focus in Mip6 role in the regulation of the HS response and the transcriptional memory defects associated to its absence, deepening in its crosstalk with transcription factors Msn2/4 and the HDAs Rpd3. Interestingly, the absence of *MIP6* provokes a differential behavior of gene expression in a set of genes (including *MSN4*), enhancing the up and down-regulation of many of them upon memory. In addition, an interesting link Mip6 and the Rpd3L complex was found in this analysis, since three genes encoding subunits of this complex were affected by memory to heat shock only in the *mip61′* mutant, which suggests a role for Mip6 in regulating the expression/degradation of these genes.

### Mip6 contribute to MSN2/4 alternative pathways upon severe stress

As shown here, *msn2Δmsn4Δmip6Δ* is the only mutant that presents a clear defect growth under severe HS. This result does not only indicate a genetic interaction of *MSN2*, *MSN4* and *MIP6* upon severe HS, but it also suggests the participation of these proteins in different cell survival mechanisms that are non-redundant upon these conditions. Mip6 regulates the expression of several Hsf1-dependent genes, since different subsets of Hsf1- regulated transcripts were either overexpressed or underexpressed in the *mip6Δ* mutant under both optimal and HS conditions^22^. Besides, Hsf1 was described to be essential for cell adaptation to a prolonged heat stress ^28^. Therefore, Mip6 may act as an Hsf1 regulator during an extended exposure of severe HS, which would explain that the lack of both Mip6 and Msn2/4 affect cell survival more than the corresponding single mutants. Further work combining mutations of all of them will help to understand better Mip6 function.

On the other hand, after an oxidative shock, differences in the growth rate of the studied mutants are observed. The lack of both Msn2/4 recovers the WT growth rate, which could be explained, as in a mild HS, by the activation of an alternative pathway that properly triggers the stress response in absence of both. However, the combination of these mutations with *MIP6* deletion leads to a growth defect, which indicates that Mip6 could positively contribute to this alternative pathway. Surprisingly, *mip6Δ* also recovers the growth problem observed in the *msn2Δ* mutant. Therefore, Mip6 may be also acting as a negative regulator of the alternative pathway. These results require further experiments to characterize specifically the role of Mip6 upon each stress.

### Is Mip6 a negative regulator of HDAC Rpd3?

Mip6 role in regulating histone acetylation levels seems to be more related with deacetylation than with the HATs, since *MIP6* genetically interacts with the HDA Rpd3 under both optimal and stress conditions, but not genetic interaction with *GCN5* was observed. Here we report that the *rpd3Δmip6Δ* double mutant does affect cell growth rate under optimal conditions and HS. This result suggests a Mip6 role in histone deacetylation, probably modulating Rpd3 function. Mip6 physically interacts with Rpd3 under both optimal and HS and the different Rpd3 complexes regulate the same target genes as Mip6 (Hsf1- and Msn2/4-dependent and RPG) ^21,22,33,36–38^. Besides, *HSP12* and *CTT1* mRNA levels decrease in the *rpd3Δmip6Δ* strain, just like in *rpd3Δ* and contrary to the phenotype of the *mip6Δ* single mutant. This suggests that Mip6 and Rpd3 may regulate the expression of these genes, participating in the same pathway, being the presence of Rpd3 essential for Mip6 function. Thus, Mip6 may act as a Rpd3 negative regulator.

Whether Mip6 interacts with Rpd3 regardless of the complex it belongs to, or Mip6 only associates with a specific Rpd3-subcomplex is not clarified yet. Interestingly, Rpd3µ gains importance during osmotic and oxidative stress and, in addition, Rpd3 is recruited to the promoters of stress response genes independently of the other Rpd3µ subunits^39,40^. It would be interesting to study a Mip6 possible participation in this recruitment, as well as Mip6 physical interaction with the other Rpd3 complexes subunits in order to deepen in the crosstalk between this two proteins. Of note, histone modifications, particularly methylation and deacetylation, appear to play a crucial role in maintaining transcriptional memory, not just in yeast but also in higher eukaryotes^16,18,35,41,42^. The presence of key histone modification-related proteins among the genes with enhanced memory and the phenotypical studies linking Mip6 to HDAC suggest that epigenetics are central to regulating heat stress transcriptional memory in yeast.

## Methods

### Microbiological and DNA recombinant techniques

*S. cerevisiae* strains and plasmids used in this work are listed in Table 1 and Table 2, respectively. For yeast growth, genes deletions and transformation, standard methods were used^21^. The created strains were checked by western blotting and/or PCR analysis. Yeast cultures were grown in liquid media at the indicated temperatures. The media used were Yeast Peptone Dextrose (YPD) or Synthetic Complete (SC). For the growth assays, cells were diluted to a OD_600_ 0.1 and 260 µL of the cells were transferred to a 96-well plate and the plated was inserted into a TECAN Spark® Multimode Microplate Reader, where the cells were incubated at 30 °C, 37 °C or 42 °C until they reached stationary phase and OD_600_ of the samples were measured every 20 min.

**Table 2.**
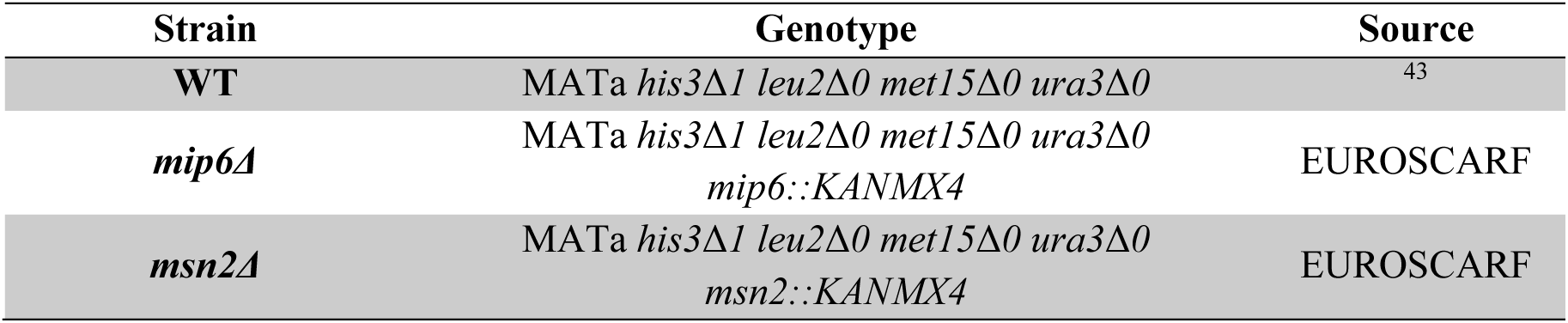
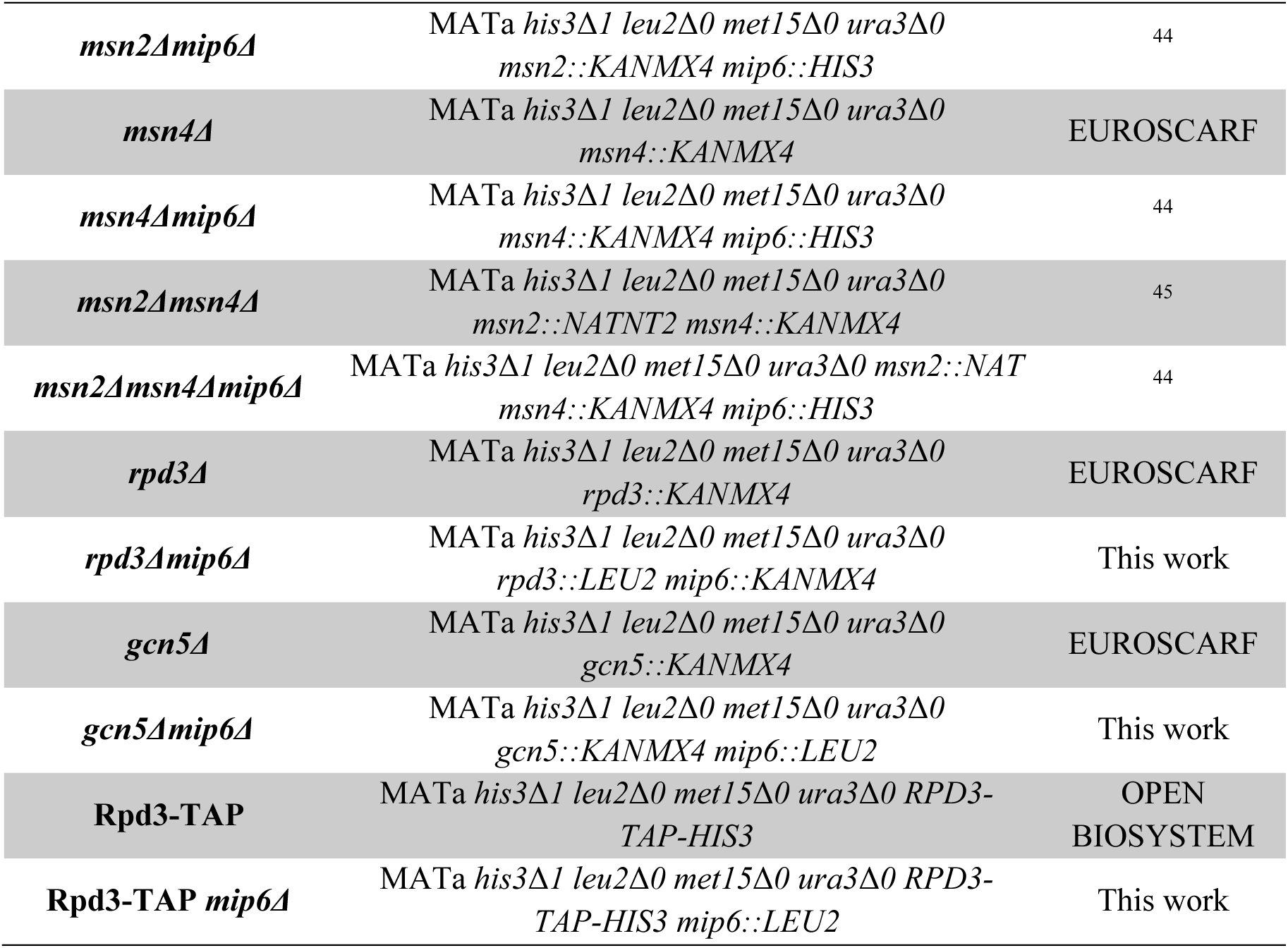
Yeast strains used during this study, with their background, their geneDc descripDon and their source.

### Stress treatments

For heat shock treatments, cells were grown at optimal conditions until exponential phase and were split into two different flasks: one for the heat shock treatment and the other one for the non-stress control. In the heat shock treatment flask, one part of the volume of the corresponding medium at 51 °C was added so that the resulting temperature of the mixture was 39 °C. Immediately, the cells were incubated at 39 °C for 20 min. In the non-stress control flask, one part of the volume of the corresponding medium at 30 °C was added.

For the oxidative stress treatment, a yeast cell survival protocol was followed^47^. Yeast cells grown in YPD until exponential phase equivalent. In the oxidative stress treatment tube, H_2_O_2_ 4 mM were added and both tubes were incubated at 30 °C for 30 min. Immediately after the incubation, cells were centrifuged and resuspended in YPD.

### Protein immunoprecipitation and western blotting

Protein immunoprecipitation was performed using GFT-TRAP® (CHROMOTEK), following the guidelines the company provide with certain modifications. Cells grown until exponential phase were resuspended in Lysis Buffer (10 mM Tris-HCl pH 7.5, 150 mM NaCl, 0.5 mM EDTA, 0.5X cOmplete™ (ROCHE), 0.5% (v/v) NP-40). Cells were broken by mixing with 0.5 mm acid-washed glass beads. The cell lysate was obtained by centrifugation at 13,000 rpm for 10 min at 4° C and incubated for 1 h at 4 °C with GFT-TRAP® beads previously equilibrated. Proteins bound to the beads were eluted with LB 4X by incubation at 95 °C for 10 min. For the western blotting, the antibodies used were α-GFP (ROCHE) and α-TAP (INVITROGEN).

### Gene expression analysis by qPCR

Total RNA was isolated by hot acid phenol extraction^48^. 10 µg of the total RNA were incubated with DNase I recombinant RNase free (ROCHE) and was then purified by phenol-chloroform extraction. cDNA was synthesized using the PrimeScript™ RT reagent Kit (TAKARA). Relative RNA levels were measured by quantitative PCR (qPCR) using the QuantStudio™ 5 Real-Time PCR System (THERMO SCIENTIFIC) or the CFX Opus 96 Real-Time PCR System (BIO-RAD). Specific pairs of primers that were used in the analysis are listed in Table 3. TB Green® Premix Ex Taq™ (Tli RNase H Plus; TAKARA) in a set volume of 10µL. The relative quantity of each PCR amplicon was measured following the 2^−ΔΔCT^ method^49^ using *SCR1* as a reference gene for normalization. Three technical replicates were performed for each sample.

**Table 3.** Yeast plasmids used during this study, with their descripDon and their source.

| Plasmid | Description | Source |
| --- | --- | --- |
| **pADH1pr-GFP** | Yeast plasmid for protein expression regulated by *ADH1* promoter and tagged in the C-terminal with a GFP epitope (*URA3* marker) | 46 |
| **pADH1pr-Mip6-GFP** | Yeast plasmid for Mip6 expression regulated by *ADH1* promoter and tagged in the C-terminal with a GFP epitope (*URA3* marker) | 44 |

**Table 4.**
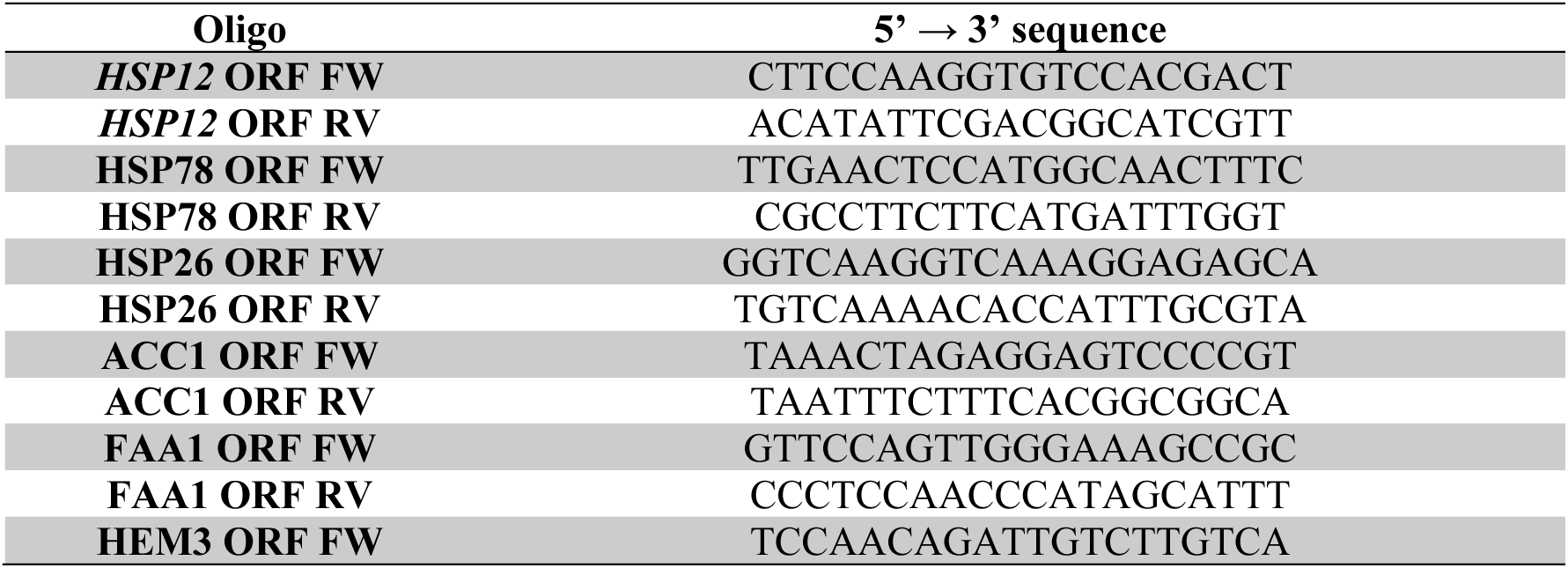
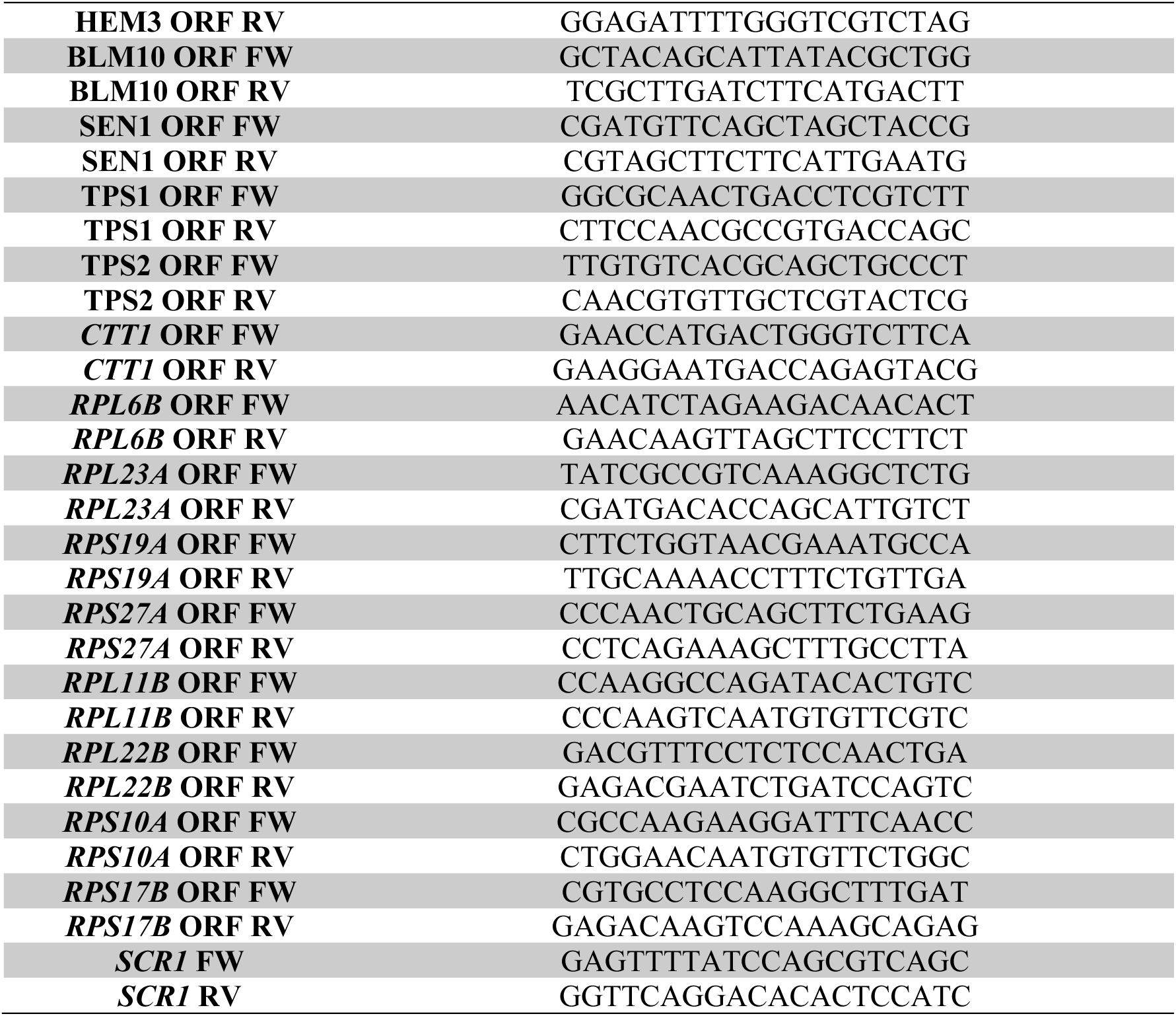
Oligos employed during this study, with their sequence in the 5’→3’ direcDon.

### Processing of mRNA datasets

For RNA-seq analyses, total RNA isolated by hot acid phenol extraction, as above, were submitted to Macrogen, Inc to be processed, using the Illumina TruSeq protocol. Between 40 and 60 million reads were obtained from each sample. The quality of the raw sequencing data was checked by fastQC and trimmomatic program was used to withdraw the adapter sequences and the bases with a low base quality from the ends. Moreover, the bases of reads which had a window size lower than 4 and a mean quality lower than 15 and the reads shorter than 36 bp were also trimmed. After trimming, quality of the samples was checked again by fastQC. Trimmed reads were mapped using sacCer3 as a reference genome with HISAT2, splice-aware aligner, and StringTie. The abundance of each gene and transcript was calculated in the read count for each sample.

The NOISeq R package^50^ was used to perform the quality control of count data. Counts were normalized via TMM^51^ and a low count filtering was applied with the NOISeq *cpm* method (with cpm = 1). Hidden batch effects were removed by ARSyN^52^. In total, we obtained gene expression values for 6,172 genes. A PCA was applied to these normalized counts to get a first glimpse over the transcriptomic changes.

### Differentially expressed genes analysis

The NOISeq method^50^ was used on the normalized counts matrix to identify differentially expressed genes in 1 to 1 comparisons. *mip6Δ* and wild-type strains at 15 and 20 min were always compared to the initial reference of the same strain. These pairwise fold changes were then represented using Sankey diagrams. Fold changes obtained by NOISeq were organised into different categories. Genes with absolute fold changes greater than 1.2 were classified as slightly differentially expressed (‘up’ or ‘down’), while those with absolute fold changes over 2 were considered strongly differentially expressed (’strong up’ or ‘strong down’). This categorization resulted in five groups. Genes showing no significant changes were labeled as ‘unchanged’.

To account for time-dependent changes following heat shock (HS), differential expression analysis was performed using the maSigPro package^53^ with default parameters.

### Statistics

For the statistical analyses, ANOVA Test, following Tukey’s Multiple Comparison Test, or t-test were performed, assuming normality and homoscedasticity of the samples, for the detection of statistically significant differences. GraphPad Prism 10 was used for the statistical analysis. In the case of genomic data, other statistics tools and R packages were used, as described above.

## Data Availability

RNA-Seq dataset is available at GEO as GSE276802. Token gpehwckyjhezhwl

## Author contributions

SR-N designed the research and supervise the work. AT-C, JS-Q and CN-C performed the experiments and analysed the genomic data. JS-Q supervised statistical methods. All authors wrote the paper. SR-N edited the manuscript.

## Supporting Information

**Table S1.** DE Genes affected by memory in the absence and the presence of Mip6 (*mip61′* and WT) related to the processes of sporulation, meiosis, and mating.

**Figure S1.** Gene expression profiles across clusters.

