## Supplementary material for "Decoding Transcriptional Memory in Yeast Heat Shock and the Functional Implication of the RNA Binding Protein Mip6": Suplemental information

***Table S1. DE Genes affected by memory in the absence and the presence of Mip6 (mip6Δ and WT) related to the processes of sporulation, meiosis, and mating.***

| | | |
|---|---|---|
| | *ACF2* | Intracellular beta-1,3-endoglucanase. Expression is induced during sporulation. |
| | *ADY3* | Protein required for spore wall formation. Subunit of leading edge protein (LEP) complex (Ssp1-Ady3-Don1-Irc10) that forms ring-like structure at leading edge of prospore membrane during meiosis II. Mediates assembly of LEP complex, formation of ring-like structure via interaction with spindle pole body components, and prospore membrane maturation. |
| | *CAC2* | Subunit of chromatin assembly factor I (CAF-1), with Rlf2p and Msi1p. Chromatin assembly by CAF-1 is important for multiple processes including silencing at telomeres, mating type loci, and rDNA. Maintenance of kinetochore structure, deactivation of the DNA damage checkpoint after DNA repair, chromatin dynamics during transcription. |
| | *CDC31* | Calcium-binding component of the spindle pole body (SPB) half-bridge. Required for SPB duplication in mitosis and meiosis II. Homolog of mammalian centrin. |
| | *CDC8* | Nucleoside monophosphate and nucleoside diphosphate kinase. Essential for mitotic and meiotic DNA replication. |
| | *CSM3* | Replication fork associated factor. Required for accurate chromosome segregation during meiosis. |
| | *DIT2* | N-formyltyrosine oxidase. Sporulation-specific microsomal enzyme involved in the production of N,N-bisformyl dityrosine required for spore wall maturation. |
| | *DNF1* | Aminophospholipid translocase (flippase). Localizes to the shmoo tip where it has a redundant role in the cellular response to mating pheromone. |
| | *FIG4* | Phosphatidylinositol 3,5-bisphosphate (PtdIns[3,5]P) phosphatase. Required for efficient mating and response to osmotic shock. |
| ***mip6Δ*** | *GIP1* | Meiosis-specific regulatory subunit of the Glc7p protein phosphatase. Regulates spore wall formation and septin organization, required for expression of some late meiotic genes and for normal localization of Glc7p. |
| | *HOP2* | Meiosis-specific protein that localizes to chromosomes. Prevents synapsis between nonhomologous chromosomes and ensures synapsis between homologs. Forms complex with Mnd1p to promote homolog pairing and meiotic double-strand break repair. Heterodimer of Hop2p-Mnd1p stimulates the Dmc1p-mediated strand invasion. |
| | *HOS2* | Histone deacetylase and subunit of Set3 and Rpd3L complexes. Required for gene activation via specific deacetylation of lysines in H3 and H4 histone tails. Subunit of the Set3 complex, a meiotic-specific repressor of sporulation specific genes that contains deacetylase activity. |
| | *HYM1* | Component of the RAM signaling network. Localizes to sites of polarized growth during budding and during the mating response. |
| | *LGE1* | Protein involved in histone H2B ubiquitination. Null mutant forms abnormally large cells, and homozygous diploid null mutant displays delayed premeiotic DNA synthesis and reduced efficiency of meiotic nuclear division. |
| | *MEI5* | Meiosis-specific protein involved in meiotic recombination. Involved in DMC1-dependent meiotic recombination. |
| | *MF(ALPHA)2* | Mating pheromone alpha-factor, made by alpha cells. Interacts with mating type a cells to induce cell cycle arrest and other responses leading to mating. |
| | *NPR3* | Subunit of the Iml1p/SEACIT complex. Null mutant has meiotic defects. |
| | *PTC1* | Type 2C protein phosphatase (PP2C). Dephosphorylates Hog1p, inactivating osmosensing MAPK cascade. Involved in Fus3p activation during pheromone response. Deletion affects precursor |

| | | |
|---|---|---|
| | | tRNA splicing, mitochondrial inheritance, and sporulation. |
| | *RAD57* | Protein that stimulates strand exchange. Involved in the recombinational repair of double-strand breaks in DNA during vegetative growth and meiosis. |
| | *REC107* | Protein involved in early stages of meiotic recombination. Involved in coordination between the initiation of recombination and the first division of meiosis. Part of a complex (Rec107p-Mei4p-Rec114p) required for ds break formation. |
| | *RIM4* | Putative RNA-binding protein. Involved in regulation of early and middle sporulation genes. Forms amyloid-like aggregates under starvation that are active in translational repression. |
| | *RSC2* | Component of the RSC chromatin remodeling complex. Required for expression of mid-late sporulation-specific genes. |
| | *SAE3* | Meiosis-specific protein involved in meiotic recombination. Involved in DMC1-dependent meiotic recombination. |
| | *SIR1* | Protein involved in silencing at mating-type loci HML and HMR. |
| | *SMT1* | Translational repressor of the mitochondrial ATP6/8 mRNA. Homozygous diploid deletion strain has a sporulation defect characterized by elevated dityrosine in the soluble fraction. |
| | *SPO20* | Meiosis-specific subunit of the t-SNARE complex. Required for prospore membrane formation during sporulation. |
| | *SPO73* | Meiosis-specific protein required for prospore membrane morphogenesis. Required for the proper shape of the prospore membrane (PSM) and for spore wall formation. Functions cooperatively with SPO71 in PSM elongation. Localizes to the PSM. Required for spore wall formation during sporulation. Dispensable for both nuclear divisions during meiosis. |
| | *BOS1* | v-SNARE (vesicle specific SNAP receptor). Required for efficient nuclear fusion during mating. |
| | *CDC55* | Regulatory subunit B of protein phosphatase 2A (PP2A). Required for correct nuclear division, chromosome segregation during achiasmate meiosis. Maintains nucleolar sequestration of Cdc14p in early meiosis. Limits formation of PP2A-Rts1p holocomplexes to ensure timely dissolution of sister chromosome cohesion. |
| | *DIG2* | MAP kinase-responsive inhibitor of the Ste12p transcription factor. Involved in the regulation of mating-specific genes and the invasive growth pathway. |
| | *DSE1* | Daughter cell-specific protein. May regulate cross-talk between the mating and filamentation pathways. Deletion affects cell separation after division and sensitivity to alpha-factor and drugs affecting the cell wall. |
| ***WT*** | *GET2* | Subunit of the GET complex. Required for the retrieval of HDEL proteins from the Golgi to the ER in an *ERD2* dependent fashion and for meiotic nuclear division. |
| | *MAD2* | Component of the spindle-assembly checkpoint complex. Delays onset of anaphase in cells with defects in mitotic spindle assembly. Regulates APC/C activity during prometaphase and metaphase of meiosis I. |
| | *MFA2* | Mating pheromone a-factor. Made by a cells. Interacts with alpha cells to induce cell cycle arrest and other responses leading to mating. |
| | *MID1* | Stretch-activated Ca2+-permeable cation channel. Required for Ca2+ influx stimulated by mating pheromones and some abiotic stresses. |
| | *MOT3* | Transcriptional repressor, activator. Role in cellular adjustment to osmotic stress including modulation of mating efficiency. Forms [MOT3+] prion under anaerobic conditions. |
| | *MSH6* | Protein required for mismatch repair in mitosis and meiosis. Forms a complex with Msh2p to repair both single-base & insertion-deletion mispairs. |
| | *MUM2* | Protein essential for meiotic DNA replication and sporulation. Subunit of the MIS complex which controls mRNA methylation |

| | |
|---|---|
| | during the induction of sporulation. |
| *NAT5* | Subunit of protein N-terminal acetyltransferase NatA. N-terminally acetylates many proteins, which influences multiple processes such as the cell cycle, heat-shock resistance, mating, sporulation, and telomeric silencing. |
| *PCH2* | Hexameric ring ATPase that remodels chromosome axis protein Hop1p. Nucleolar component of the pachytene checkpoint, which prevents chromosome segregation when recombination and chromosome synapsis are defective. Also represses meiotic interhomolog recombination in rDNA. Required for meiotic double-stranded break formation. |
| *PDS5* | Cohesion maintenance factor. Involved in sister chromatid condensation and cohesion. Also required during meiosis. |
| *PFS1* | Sporulation protein required for prospore membrane formation. Required for prospore membrane formation at selected spindle poles. Ensures functionality of all four spindle pole bodies during meiosis II. |
| *PRM1* | Pheromone-regulated multispanning membrane protein. Involved in membrane fusion during mating. Localizes to the shmoo tip. |
| *RAM1* | Beta subunit of the CAAX farnesyltransferase (FTase). This complex prenylates the a-factor mating pheromone and Ras proteins. Required for the membrane localization of Ras proteins and a-factor. |
| *RAS2* | GTP-binding protein. Regulates nitrogen starvation response, sporulation, and filamentous growth. Homolog of mammalian Ras proto-oncogenes. |
| *RCE1* | Type II CAAX prenyl protease. Involved in the proteolysis and maturation of Ras and the a-factor mating pheromone. |
| *REV3* | Catalytic subunit of DNA polymerase zeta. Involved in translesion synthesis during post-replication repair. May be involved in meiosis. |
| *RLF2* | Largest subunit (p90) of the Chromatin Assembly Complex (CAF-1). Important for mating type loci and rDNA. |
| *SCW10* | Cell wall protein with similarity to glucanases. May play a role in conjugation during mating based on mutant phenotype and its regulation by Ste12p. |
| *SCW4* | Cell wall protein with similarity to glucanases. scw4 scw10 double mutants exhibit defects in mating. |
| *SLK19* | Kinetochore-associated protein. Required for normal segregation of chromosomes in meiosis and mitosis. Component of the FEAR regulatory network, which promotes Cdc14p release from the nucleolus during anaphase. Potential Cdc28p substrate. |
| *SPP1* | Subunit of COMPASS (Set1C), which methylates histone H3 on lysine 4. Promotes meiotic DSB formation by interacting with H3K4me3 and Rec107p, a protein required for Spo11p-catalyzed DSB formation located on chromosome axes. |
| *STE18* | G protein gamma subunit. Forms a dimer with Ste4p to activate the mating signaling pathway. |
| *STH1* | ATPase component of the RSC chromatin remodeling complex. Required for expression of early meiotic genes. Promotes base excision repair in chromatin. |
| *SUM1* | Transcriptional repressor that regulates middle-sporulation genes. Required for mitotic repression of middle sporulation-specific genes. Regulated by pachytene checkpoint. |
| *SUR7* | Plasma membrane protein, component of eisosomes. Sporulation and plasma membrane sphingolipid content are altered in mutants. |
| *TEP1* | PTEN homolog with no demonstrated inositol lipid phosphatase activity. Plays a role in normal sporulation. |

***Figure S2. Gene expression profiles across clusters. Heatmaps displaying the gene expression levels of all genes within the four distinct clusters.*** *The data represent biological replicates for both WT and* mip6Δ *strains, measured at three different HS time points, under memory and no-memory conditions. Each cluster highlights genes with shared expression dynamics across the experimental conditions.*

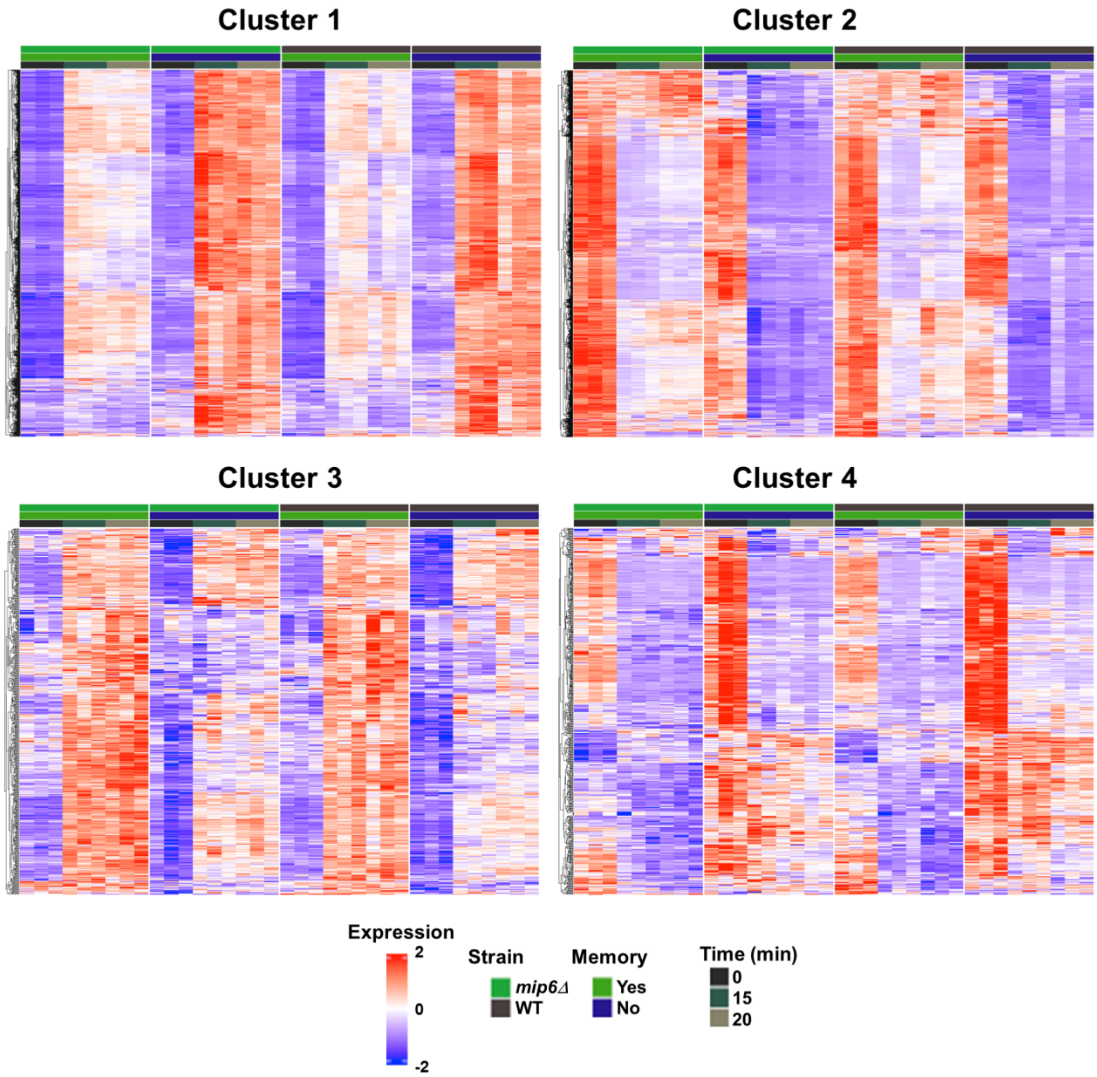
